# Pili allow dominant marine cyanobacteria to avoid sinking and evade predation

**DOI:** 10.1101/2020.07.09.194837

**Authors:** Maria del Mar Aguilo-Ferretjans, Rafael Bosch, Richard J. Puxty, Mira Latva, Vinko Zadjelovic, Audam Chhun, Despoina Sousoni, Marco Polin, David J. Scanlan, Joseph A. Christie-Oleza

**Affiliations:** University of the Balearic Islands, Palma, Spain; IMEDEA (CSIC-UIB), Esporles, Spain; School of Life Sciences, University of Warwick, Coventry, UK; Department of Physics, University of Warwick, Coventry, UK

## Abstract

How oligotrophic marine cyanobacteria position themselves in the water column is currently unknown. The current paradigm is that these organisms avoid sinking due to their reduced size and passive drift within currents. Here, we show that one in four picocyanobacteria encode a type IV pilus which allows these organisms to increase drag and remain suspended at optimal positions in the water column, as well as evade predation by grazers. The evolution of this sophisticated floatation mechanism in these purely planktonic streamlined microorganisms has profound implications for our current understanding of microbial distribution in the oceans, predator-prey interactions and, ultimately, will influence future models of carbon flux dynamics in the oceans.

---

A quarter of all primary production on Earth occurs in large nutrient deplete oceanic gyres^1^. Primary production in these large biomes is mainly driven by the dominant marine cyanobacteria *i.e. Prochlorococcus* and *Synechococcus*^2^. Gyres are permanently thermally stratified, where a lack of upward physical mixing poses a challenge for the microbial communities that inhabit them. How then do purely planktonic cyanobacterial cells in suspension combat the downward pull of gravity through the biological pump –*i.e.* drawing fixed carbon towards the ocean interior? Moreover, how do these highly-specialised planktonic microbes place themselves in their ‘preferred spot’, such as the well-established vertical distribution of high and low light-adapted *Prochlorococcus* ecotypes^3–5^? Marine picocyanobacteria lack flagellar structures for swimming or gas vacuoles for flotation^6^, and only a limited number of *Synechococcus* strains possess non-conventional mechanisms for motility^7,8^. Therefore, it has been assumed that these free-living microbes avoid sinking due to their lower density and reduced size^9^, and bloom when they encounter their optimal environmental conditions while drifting randomly within marine currents.

To date, type IV pili are known to provide functions such as twitching motility, surface attachment, biofilm formation, pathogenicity, as well as conjugation, exogenous DNA acquisition and competence^10–12^. These extracellular appendages can be rapidly extended and retracted by polymerising and depolymerising cycles of the major pilin subunit *e.g.* PilA, requiring a defined transmembrane apparatus and the consumption of energy in the form of ATP^11,13^. Functional analysis of most type IV pili has focused on pathogenic microbes and their use of surfaces or substrates for pilus anchoring. However, analysis of pili from freshwater cyanobacteria has revealed these appendages can be used for twitching motility during phototaxis in *Synechocystis*^14^ or exogenous DNA acquisition in both *Synechocystis* and *Synechococcus elongatus*^15,16^. This latter function requires an additional set of proteins for competence such as ComEA and ComEC. While acquisition of DNA by *S. elongatus* is performed by the third of three PilA-like proteins encoded by this strain (PilA3)^16^, no known function has been attributed to PilA1 and PilA2 other than being dispensable for attachment and biofilm formation^17^. A mutant in *S. elongatus* that no longer produced PilB –the protein responsible for pilus elongation– abolished the production of pili appendages, made up of PilA1, and was reported to suppress planktonic growth of this strain^17^. The presence of pilus genes in purely planktonic marine microbes has previously been reported, but their role remains enigmatic^18^.

Here, we show that almost a quarter of all marine picocyanobacteria encode a PilA1-like pilus. We show that this extracellular appendage produced in these purely planktonic organisms –which rarely encounter any kind of surface in their natural habitat– allows cells to increase drag and remain in an optimal position in the water column as well as avoid being preyed upon. This provides yet another biological function to these filamentous appendages and sheds light on the ecological role of type IV pili in marine ecosystems.

## RESULTS AND DISCUSSION

### Abundant production of a type IV pilus in *Synechococcus* sp. WH7803

We first detected an abundant PilA protein (*i.e.* SynWH7803_1795) in the extracellular proteomes of the model marine cyanobacterium *Synechococcus* sp. WH7803, accounting for up to 25% of the exoproteome^19,20^. Transmission electron microscopy confirmed the existence of the macromolecular pili structures (Fig 1A and Fig S1). Unlike *Synechocystis* sp. PCC6803 that simultaneously produces thick and thin pili^14^, this marine picocyanobacterium presented multiple pili of similar thickness, each ~10 μm in length. The amino acid sequence of PilA revealed a typical sec-targeting signal peptide and a conserved GFTLxE motif at the N-terminus of the protein (Fig 1B) that is known to be cleaved in the cytoplasmic membrane by PilD before the protein is translocated to the base of the pili for assembly^10^. After cleavage, the N-terminal of PilA can be post-translationally modified, *e.g.* methylated, to increase the hydrophobicity and stability of the pilin^10,21^, although we were unable to detect this modified N-terminus tryptic peptide during proteomic analyses. In close proximity to *pilA* in the *Synechococcus* sp. WH7803 genome we found five other type IV-like pilin genes (Fig 1C), all with the conserved GFTLxE motif (Fig 1D).

**Fig 1.**
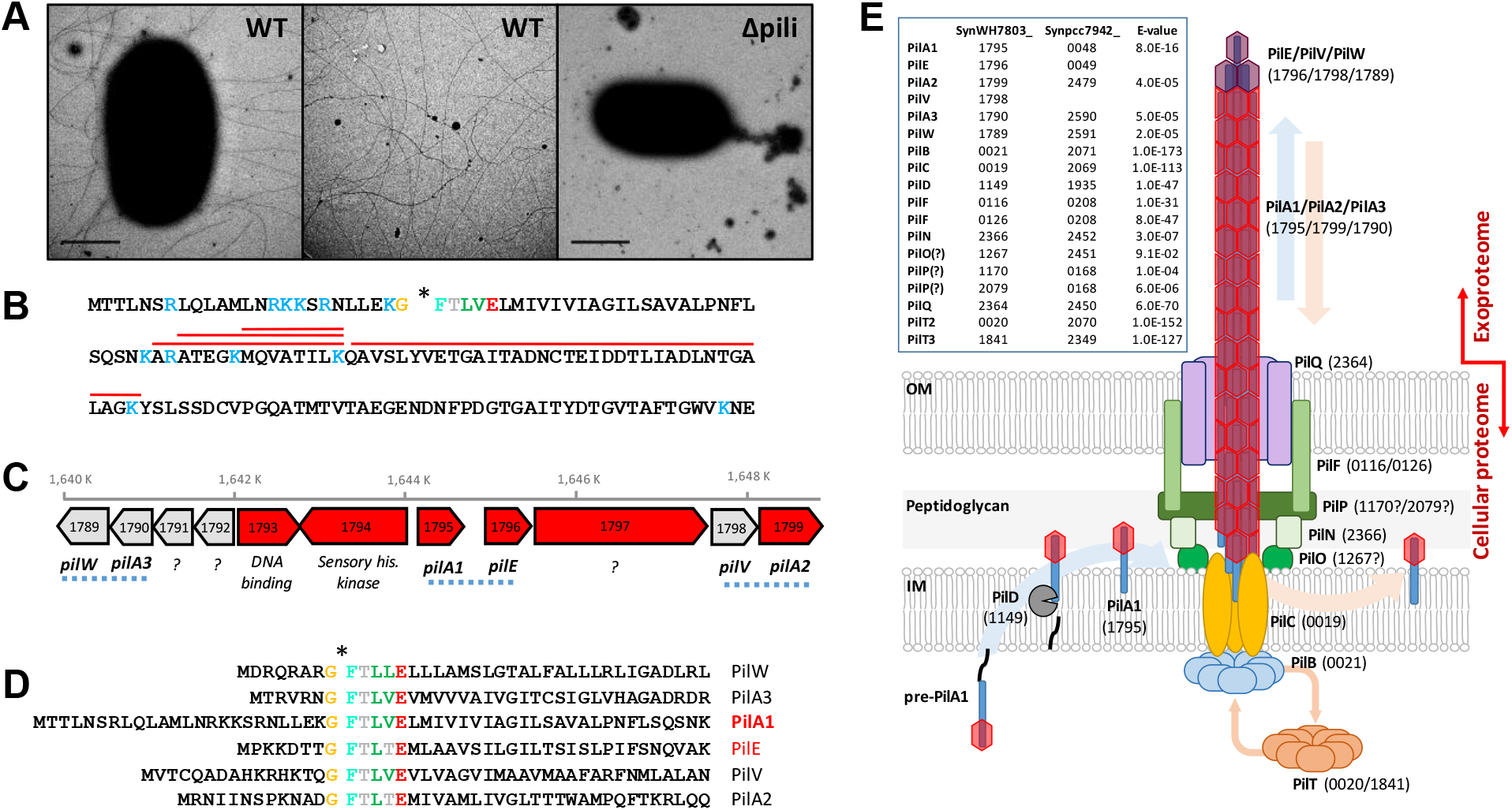
Pilus in the marine cyanobacterium *Synechococcus* sp. WH7803. **(A)** Transmission electron microscopy images of wild type *Synechococcus* sp. WH7803 (WT) and pili mutant (Δpili) obtained from late exponential liquid cultures incubated in ASW medium under optimal growth conditions. Long pili appendages were only observed in the wild type strain (Fig S1). Middle panel image, obtained with the same magnification as other panels, is from an intercellular region between wild type cells to improve the visualisation of the pili. Scale bar represents 1 μm. **(B)** The amino acid sequence of PilA1 (SynWH7803_1795). Trypsin hydrolytic sites are indicated in blue. Red lines highlight tryptic peptides detected by shotgun proteomics. The conserved GFTLxE motif is shown and the cleavage site is indicated with an asterisk. **(C)** Genomic context of *pilA1* in *Synechococcus* sp. WH7803. Numbers in each gene represent their ID number (SynWH7803_). In red are genes detected by proteomics. While PilA1 and PilE are abundantly detected in exoproteomes^20^, PilA2 has only ever been detected in cellular proteomes of this strain^22^. Blue dotted lines indicate genes encoding possible structural pilin pairs, *i.e.* PilA1-PilE, PilA2-PilV and PilA3-PilW. Question marks indicate genes encoding proteins of unknown function. **(D)** The N-terminal amino acid sequence of PilA1 and five other pilin-like proteins, all with the highly conserved GFTLxE motif. **(E)** *Synechococcus* sp. WH7803 structural pilus proteins identified by homology with *S. elongatus* PCC 7942^16^ and assembled as modelled by Craig *et al*^11^.

Using the pilus apparatus from the freshwater cyanobacterium *S. elongatus* PCC 7942 as a reference^16^ and the established architecture for type IV pilus machinery^11^, we were able to find all components necessary for pilus assembly in *Synechococcus* sp. WH7803 (Fig 1E). The genetic cluster encoding the six pilin-like proteins (Fig 1C and 1D) likely produces three types of pili that use the same transmembrane pilus structure. Based on homology with the annotated genes from *S. elongatus*^16^ and conserved domains found using the CD-search tool in NCBI, we suggest the three pili types: PilA1-PilE, PilA2-PilV and PilA3-PilW (Fig 1C). Of these types, shotgun proteomic analyses have only ever detected PilA1-PilE^19,20^, although PilA2 was also detected in low abundance in cellular –but not extracellular– proteomic datasets^22^. Unlike in *S. elongatus*, where PilA1 and the contiguously-encoded pilin-like protein are almost identical, the amino acid sequence of PilA1 and PilE in *Synechococcus* sp. WH7803 are clearly distinguishable. Although in much lower abundance, PilE seems to be correlated with PilA1 in the exoproteomes of this cyanobacterium^19,20^ and, therefore, it is possible that PilE and PilA1 form subunits of the same pilus apparatus.

### Pilus distribution amongst picocyanobacterial isolates and Single-cell Assembled Genomes (SAGs)

Genomic analysis of sequenced marine picocyanobacterial isolates downloaded from the Cyanorak database revealed that 74% of sequenced *Synechococcus* (n=46) and 33% of *Prochlorococcus* (n=43) encoded *pilA1* (Fig 2 and Table S1). In *Synechococcus*, the pilus was prevalent in all clades (93%; n=28) except for clades II and III where it was less abundant (44%; n=18). Interestingly, all low light *Prochlorococcus* isolates from clades III and IV encoded *pilA1* (n=7; Fig 2). Most of these *pilA1*-containing strains also encoded a *pilE* homologue in close proximity. Genes *pilA2* and *pilA3* were also abundantly found in *Synechococcus* (59 and 74%, respectively), although were much less prevalent in *Prochlorococcus* (12 and 9%, respectively). As expected, all strains that encode at least one of the *pilA* types also possessed the transmembrane pilus apparatus, whereas this apparatus was completely absent or partially lost in strains lacking *pilA* (Fig 2). PilA3 is known to be involved in DNA uptake and competence in *S. elongatus*, requiring additional competence proteins to do so^16^. Marine picyanobacteria are not known for being naturally competent but, interestingly, all strains encoding PilA3 also contained the competence genes encoding ComEA and ComEC (Fig 2). Further work is needed to investigate the conditions under which the PilA3-type pilus becomes active in these organisms and, therefore, when exogenous DNA might be taken up.

**Fig 2.**
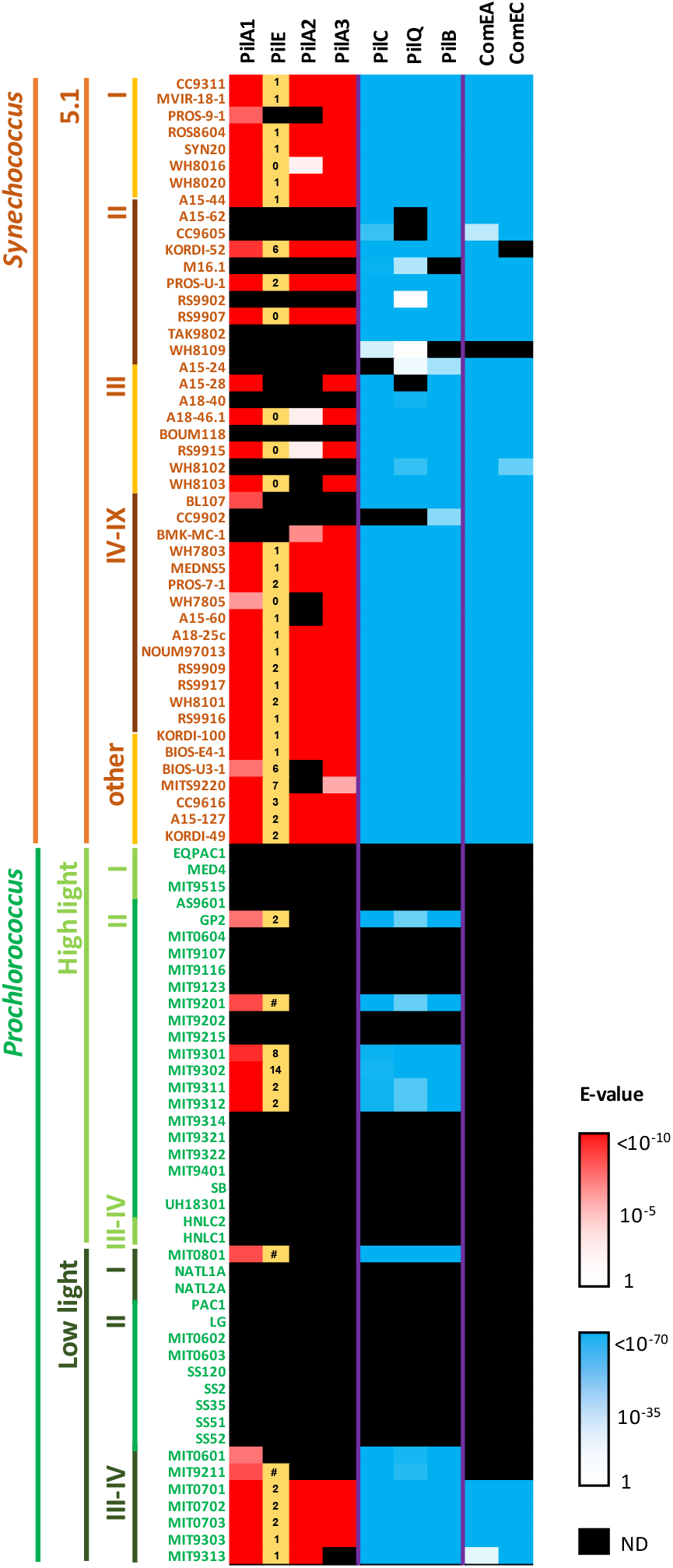
The presence of pilus-related proteins in cultured marine picocyanobacteria strains. Pilus proteins from *Synechococcus* sp. WH7803 were used for the BLASTp search. Log10 E-value scales are shown (1 to <10^−10^, white to red; and 1 to <10^−70^, white to blue). Black cells represent proteins that were not detected (ND). Numbers in the ‘PilE’ column indicate the genomic distance between *pilE* and *pilA1* homologues (*e.g.* 1 denotes *pilE* and *pilA1* are contiguous in the genome; 0 denotes the same gene gave homology to both *pilE* and *pilA1* due to the conserved N-terminal of the protein; # denotes both genes are separated by >20 genes). PilC, PilQ and PilB were used to determine the presence of the pilus transmembrane apparatus. ComEA and ComEC were selected to determine the presence of the additional machinery required for competence.

The GFTLxE motif is conserved in 87.5% of PilA1 sequences encoded by cultured marine picocyanobacteria (*i.e.* 42 of the 48 sequences; Table S1). The remaining six PilA1 sequences possess a GFSLxE motif, five of which were in *Prochlorococcus* strains. Across the full length of the mature PilA1 protein, which on average is 140 amino acids long, only the first ~50 N-terminal amino acids starting from the conserved GFTLxE motif are well conserved amongst all sequences, a commonly observed feature in PilA-like proteins^11^. The C-terminus of the protein showed a remarkably high variability even between closely related strains. Despite this high variability, their predicted structures were still similar to those of known pili subunits (Fig S2)^23,24^. During pili assembly, the helix encoded by the conserved N-terminus of PilA remains in the pilus core and only the variable C-terminus –that producing anti-parallel β-sheets– is exposed to the milieu^11^. We hypothesise that the hyper-variability of the exposed C-terminus is a strategy to escape phage attachment, it being a known pathway used by phage for host encounter and infection^25,26^. Similarly, flagella have a hyper-variable region, which has also been attributed to phage and immune system evasion^27^.

The screening of 190 picocyanobacterial Single-cell Assembled Genomes (SAGs) obtained from surface seawater across the globe^28^ –those with over 75% completeness– revealed the presence of *pilA1* and genes encoding components of the pilus apparatus in almost one in four marine picocyanobacteria (24% encoded *pilA1* and 21-25% the pilus apparatus; Table 1 and Table S1). As expected by its abundance in the oceans, *Prochlorococcus* comprised almost 96% of all 190 SAGs (Table 1). The prevalence of the pilus was much higher amongst the low light *Prochlorococcus* SAGs from clades II/III (67-80%) than in those belonging to other ecotypes, and *Synechococcus* showed the abundance observed in cultures isolates (~76%; Table 1).

**Table 1.**
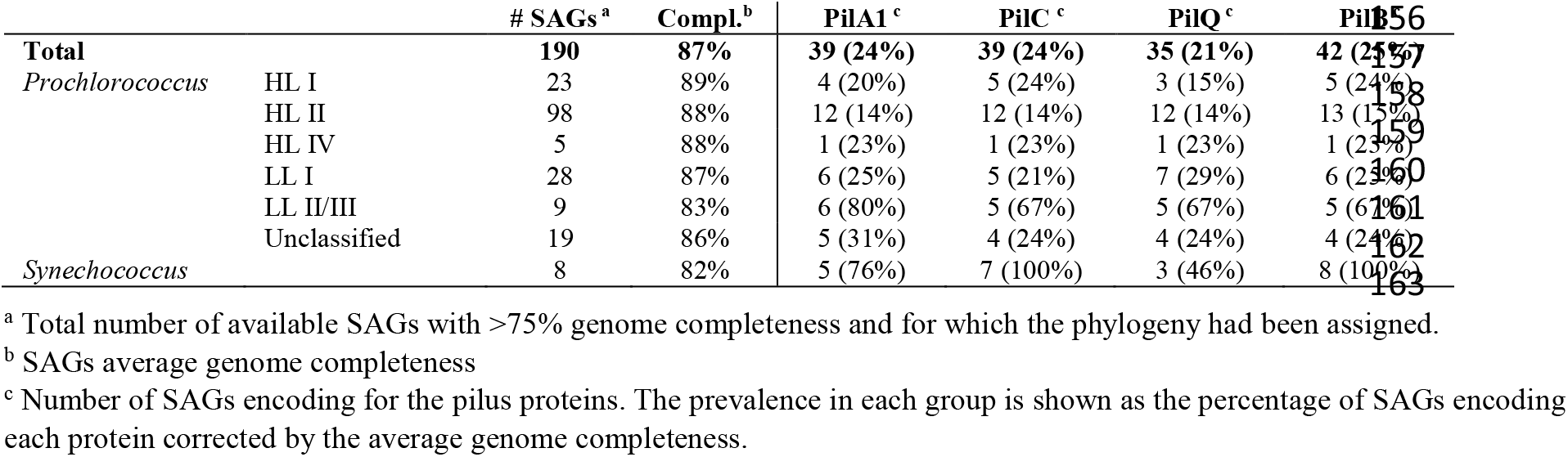
PilA1 and pilus apparatus distribution amongst planktonic marine SAGs.

### Global distribution and expression of picocyanobacterial *pilA1* in marine pelagic ecosystems

The distribution and expression of *pilA1* in the surface ocean was determined by analysing its presence in the global marine TARA metagenome and metatranscriptome datasets. An HMM profile generated from the PilA1 sequences (cultured isolates and SAGs; Table S1) was used to search the TARA datasets in the Ocean Gene Atlas portal^29^ retrieving 903 and 837 individual hits from the metagenomes and metatranscriptomes, respectively (using a cut-off E-value < 10^−10^). Sequences assigned to *Prochlorococcus* represented 85% in both datasets, whereas those assigned to *Synechococcus* represented 12% and 11% of the metagenomes and metatranscriptomes, respectively. BLAST analysis of these hits against PilA1, PilA2 and PilA3 sequences was used to confirm the specificity of our HMM profile, proving effective in discriminating against PilA2 and PilA3 sequences (each representing less than 1% of the hits).

The abundance and transcription of genes encoding PilA1 from *Prochlorococcus* and *Synechococcus* across all oceanic regions, marine biomes and water depths (Fig 3) revealed a similar abundance of *pilA1* from *Prochlorococcus* in both metagenomic and metatranscriptomic datasets. In contrast, *pilA1* from *Synechococcus* was enriched in the metatranscriptomes, mainly driven by the increased expression in the North Atlantic Ocean and Mediterranean Sea as well as in coastal and westerlies, biomes where *Synechococcus* are known to thrive. As expected, a large reduction in the presence and expression of the picocyanobacterial *pilA1* was noted in the polar oceans, where both genus are not abundant. Furthermore, the presence and transcription of cyanobacterial *pilA1* decreased drastically in the aphotic mesopelagic layer, in accordance with picocyanobacteria naturally populating only euphotic layers of the ocean.

**Fig 3.**
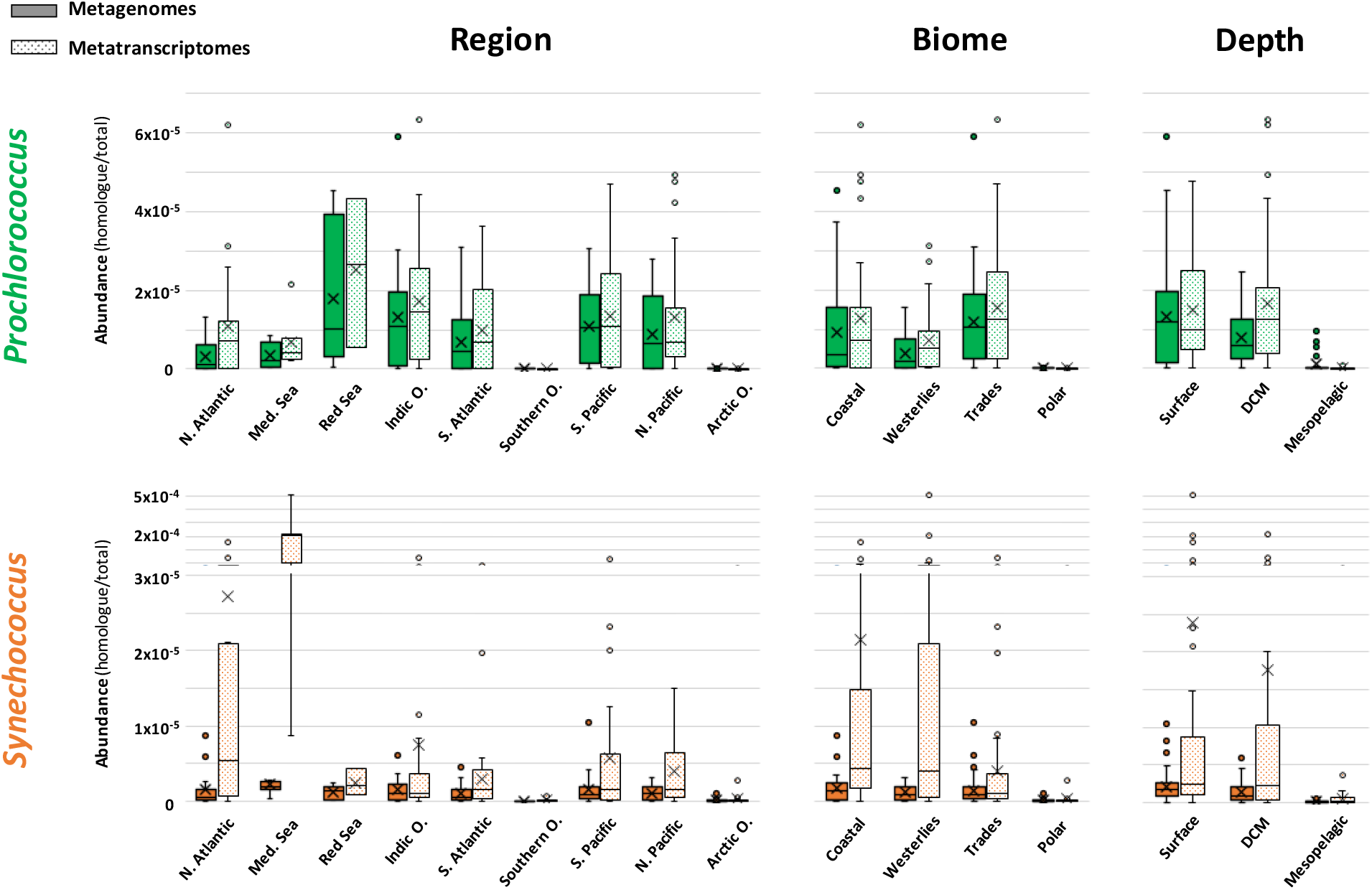
Distribution of PilA1 amongst marine ecosystems. Whisker box plots showing *pilA1* gene abundance in *Prochlorococcus* and *Synechococcus* in metagenomes and metatranscriptomes generated from all filters (0.2-3 μm) and sampling stations of the TARA Oceans global marine survey. Abundance was calculated by dividing the sum of the abundances of *pilA1* homologs assigned to each genus by the sum of total gene abundance from all reads from the sample. Data is presented by oceanic region, biome and water depth *i.e.* surface, deep chlorophyll maxima (DCM) and mesopelagic layers. Filters from polar regions were excluded from the depth abundance analysis. Exclusive median (line), average (cross) and atypical values (circles) are also indicated.

### PilA1-type pili increase drag and allow cells to remain planktonic

The extensive distribution and expression of such an extracellular appendage begs the question why such a complex structure is so prevalent in such streamlined planktonic cyanobacteria. To assess this, and to assign a biological function to this extracellular structure in the context of the ecology of marine planktonic bacteria more generally, we abolished the production of PilA1 and PilE in *Synechococcus* sp. WH7803. As expected, the fully segregated pili mutant strain no longer produced the extracellular structure (Fig 1A and Fig S1). Most remarkably, we observed a clear loss in the strain’s ability to remain suspended in its typical planktonic form (Fig 4A). By tracking wild type and pili mutant cells of *Synechococcus* sp. WH7803, we determined that the lack of pili produced an average cell sinking rate of 8.4 ± 0.4 mm/day, while the wild type had an average uplifting drift of 0.8 ± 0.3 mm/day (Fig 4B, Suppl. Movies 1-4 and full data in Table S2). While avoiding sedimentation, the pili did not appear to confer motility, with both mutant and wild type strains producing ‘pin-prick’ colonies in sloppy agar plates as opposed to fuzzy colonies characteristic of motile strains^8^. Furthermore, neither the mutant nor the wild type strain aggregated when grown in shaken liquid cultures.

**Fig 4.**
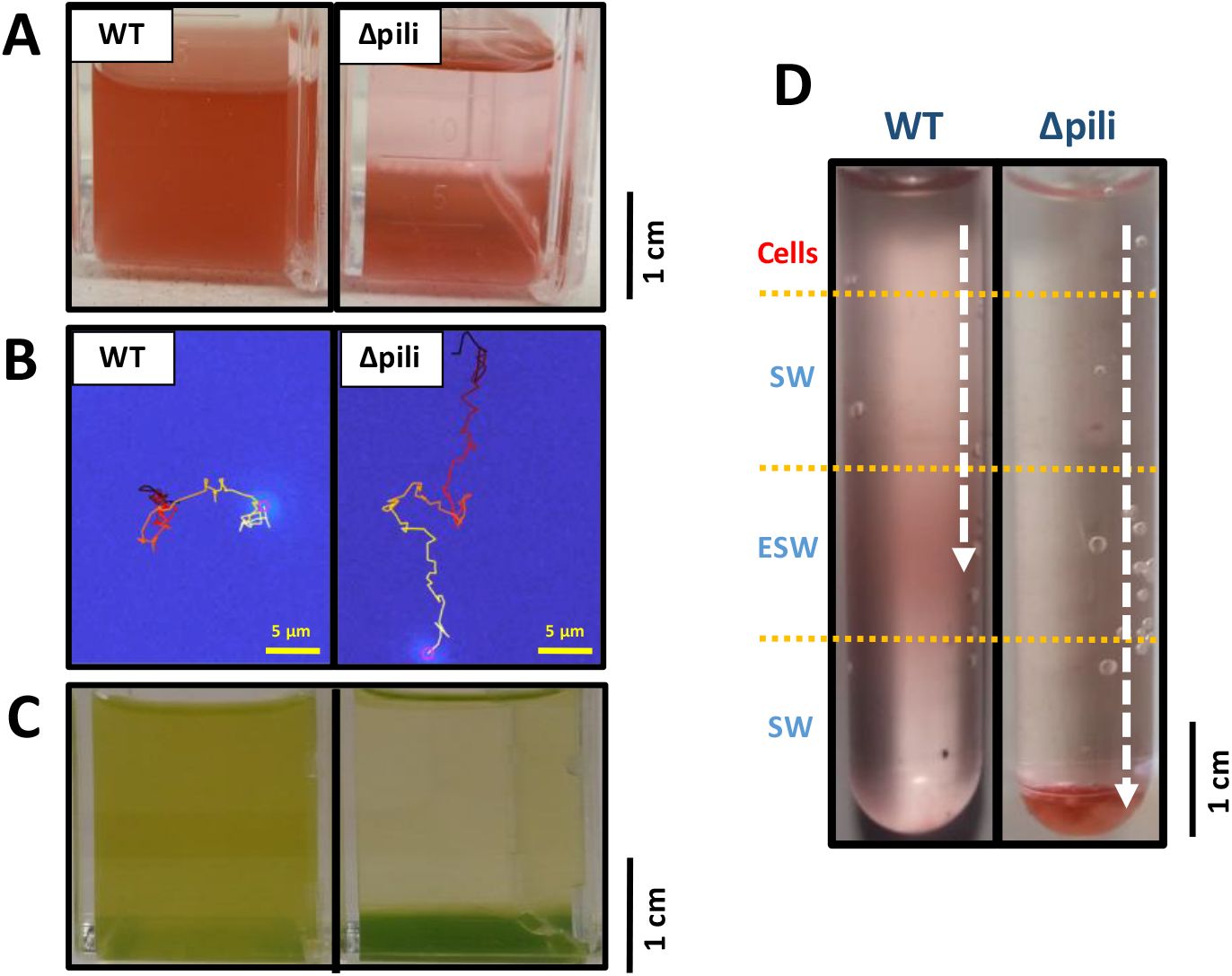
Pili avoid sinking in planktonic marine picocyanobacteria. **(A)** Non-shaken cultures of wild type (WT) and pili mutant (Δpili) of *Synechococcus* sp. WH7803. The pili mutation and phenotype has proven very stable with the mutant strain unable to remain in a planktonic form when not shaken. The appearance of the mutant in the image is consistently achieved after 2-3 days of no shaking. **(B)** Example of tracked suspended cell from a wild type and pili mutant culture of *Synechococcus* sp. WH7803. The movement was tracked over 100 s and movement is indicated from dark red (start) to yellow (end). Full data of all tracked cells can be found in Table S2 and examples in Movies S1-S4. **(C)** Non-shaken *Prochlorococcus* sp. MIT9313 cultures grown in Pro99 medium (left) and PCR-S11 medium (right). The appearance of cells in the image is consistently achieved after 2-3 days of no shaking. PilA1 (PMT_0263) was only detected in the exoproteome of suspended cultures (left; Table S4). **(D)** Nutrient step gradient column where wild type and pili mutant cells of *Synechococcus* sp. WH7803 were placed at the top. Nutrient deplete (SW) and nutrient enriched layers (ESW) are indicated. Cells were harvested by centrifugation from late-exponential cultures grown under optimal conditions, and resuspended in SW. The arrow indicates the trajectory made by the cells, and where they accumulated after three days. Other gradients tested are shown in Fig S4. Images represent one of three culture replicates.

We performed a comparative proteomic analysis between wild type *Synechococcus* sp. WH7803 and the pili mutant to assess any additional effects of disrupting the PilA1-PilE pilus (Table S3). Apart from the complete absence of the PilA1 and PilE proteins, only three other proteins were significantly down-regulated in the cellular proteome of the pili mutant strain (Fig S3): SynWH7803_0049 (12.9 fold) and SynWH7803_1797 (3.2 fold), both of unknown function and, most interestingly the pilus retraction protein PilT (SynWH7803_0020; 2.9 fold reduction). SynWH7803_1797 is located just downstream of PilA1-PilE and is predicted to encode a secreted protein that is usually found in low abundance in the exoproteome of *Synechococcus* sp. WH7803^20^. Indeed, it was also found down-regulated in the exoproteome of the pili mutant (2.5x; Fig S3). The exoproteomes also revealed a generalised shift of proteins more abundantly detected in the pili mutant. These were mainly low abundance cytoplasmic proteins that were barely detected in the wild type strain (Fig S3 and Table S3). Most likely, the absence of the abundant PilA1 protein from the exoproteome of the mutant strain caused an artifactual increase in the detection of lower abundance proteins by mass spectrometry and, despite efforts to normalise the data, there was an apparent upregulation of most proteins.

We had previously observed that *Prochlorococcus* sp. MIT9313 (a strain that encodes PilA1; Fig 2) routinely switches between planktonic and sedimenting lifestyles when grown in different media (Pro99 and PCR-S11, respectively; Fig 4C). Whilst *Prochlorococcus* remains genetically intractable^30^, we compared the exoproteomes of this strain in both medias. Commensurate with our findings in *Synechococcus* sp. WH7803, PilA1 was not detected in the exoproteomes of sedimenting *Prochlorococcus* sp. MIT9313 cultures, whereas it was present in all of the planktonic ones (*i.e.* PMT_0263; Table S4), suggesting *Prochlorococcus* can coordinate production of the pili in response to the distinct nutrient environments present in both medias.

### PilA1 allows cells to retain an optimal position within the water column

We further explored the effect of nutrient stress on pili production in *Synechococcus* sp. WH7803, with nitrogen and metal depletion showing the strongest decline (*i.e.* 11.5 and 3.4-fold decrease in PilA1 production, respectively), whilst phosphorus depletion had no effect (Table S5). Although in low abundance, PilA1 of *Synechococcus* sp. WH7803 was detected even under natural oligotrophic conditions (*i.e.* when incubated in natural seawater; representing 0.3% of the exoproteome) and showed a slight increase after adding environmentally-relevant concentrations of nutrients (0.5% of the exoproteome in the presence of 8.8 μM N and 0.18 μM P). Pili detection became most obvious under higher nutrient conditions (60-fold increase in pili at 88 μM N and 1.8 μM P when compared to nutrient deplete seawater; Table S6). Considering the biological significance of this phenotype in the context of marine planktonic organisms –where the production of pili reduces sedimentation by increasing the viscous drag of the cell– the extension/retraction of pili would allow a cell to position itself at an optimal position in the water column, *e.g.* in patches of high nutrient availability. To further investigate this, we set up a nutrient step gradient. While the pili mutant, as expected, sank through the gradient independently of nutrient availability, the wild type strain was able to position itself in the nutrient-replete layer where it remained via the production of pili (Fig 4D and Fig S4).

*Synechococcus* sp. WH7803 also retained pili over a 24 h period of darkness. Nevertheless, after three days under dark conditions –during which cultures are known to remain viable^31^, the detection of pili dropped drastically (*i.e.* >20-fold drop in pili abundance in the exoproteome; Table S7). Marine cyanobacteria occupying photic layers of the ocean will therefore keep their pili structures over normal diurnal light-dark cycles to maintain their position, but cells will cease to retain their position once they sink out of the euphotic zone, retracting their pili possibly as a strategy to recover energy while awaiting an uplift back to photic layers by upwelling currents.

### Pili prevent grazing

Given the cell surface nature of this pilus, we also assessed whether it could mediate ecological interactions with other organisms. We found that as well as preventing sedimentation, pili allow bacteria to evade predation by protist grazers. Thus, *Synechococcus* sp. WH7803 and the pili mutant grown in the presence of two bacterivorous protozoa *i.e.* the preying ciliate *Uronema* and suspension feeding flagellate *Cafeteria*, strikingly showed that whilst the wild type strain was able to completely evade grazing by *Cafeteria* and largely delay culture depletion by *Uronema*, the pili mutant was efficiently grazed by both protists (Fig 5). Presumably, the long pili appendages interfere with the way bacterivorous protozoa access their prey and, hence, this reduces their susceptibility to grazing.

**Fig 5.**
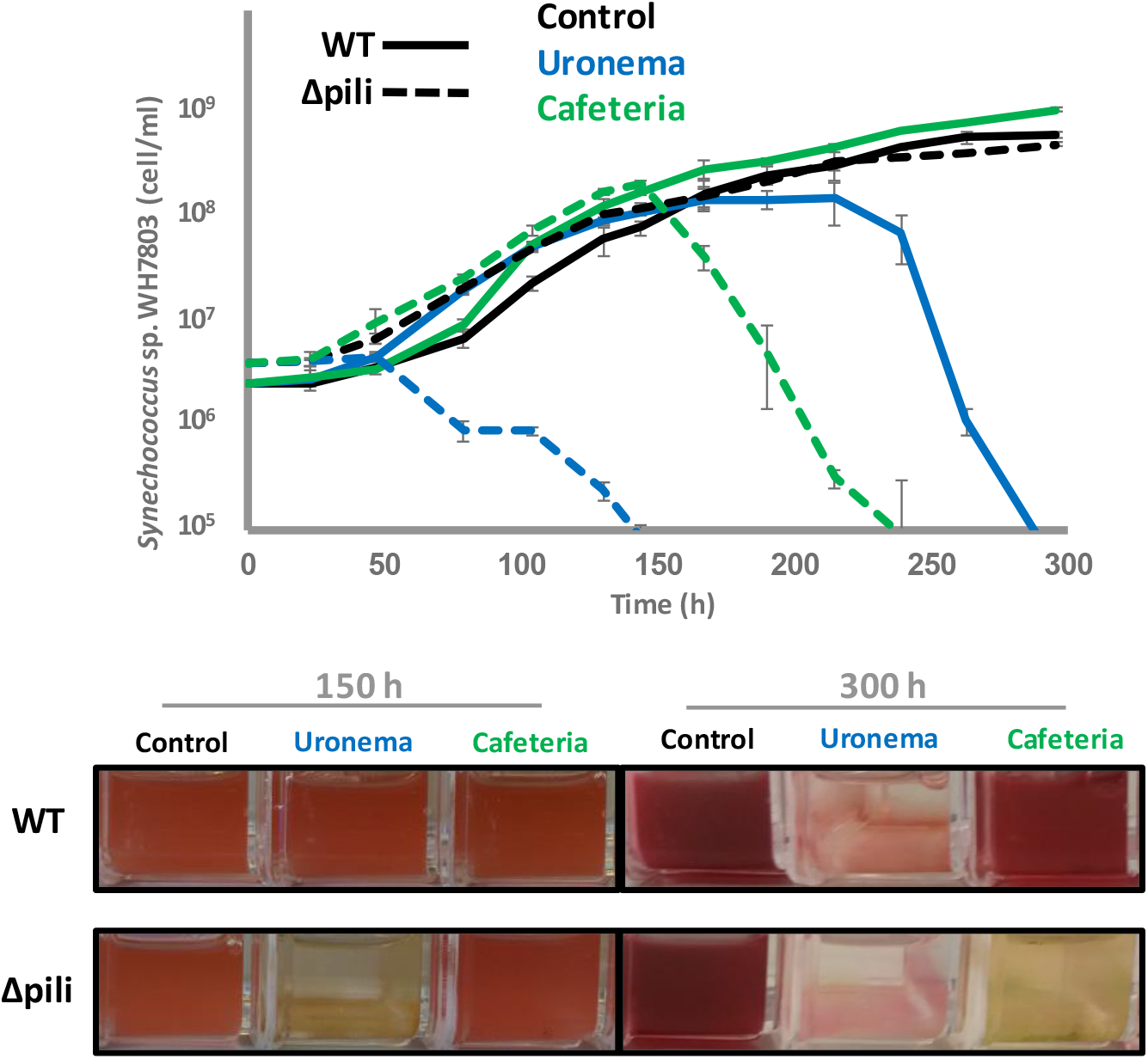
Pili confer resistance to grazing. Growth curves (top panel) and culture images (bottom panel) of wild type (WT) and pili mutant (Δpili) cultures of *Synechococcus* sp. WH7803 incubated in the absence (control) and presence of two different grazers. Cultures were subjected to constant shaking to keep the wild type and PilA1 mutant in planktonic form. Error bars represent the standard deviation from three culture replicates. Images represent one of three culture replicates.

### Concluding remarks

Together, our data suggests a novel and sophisticated mechanism that enables oligotrophic marine picocyanobacteria to stay buoyant and reduce cell death due to grazing through the use of pili, adding a new biological function to these extracellular appendages. This PilA1-type pilus, widely distributed and expressed amongst dominant marine oligotrophic cyanobacteria, allow these purely planktonic organisms to increase cell drag and, consequently, maintain an optimal position in the water column. This mechanism is under tight regulation in response to discrete stimuli *e.g.* nutrients, which may vary depending on the ecological adaptation and preferred niche of each individual organism. Therefore, as opposed to flagellated bacteria that show positive chemotaxis towards nutrient hotspots in the oceans^32^, non-motile picocyanobacteria may apply a more passive strategy which consists of elongating their pili when they encounter preferable conditions to remain in an optimal position while drifting in a water body. These long appendages also interfere with the access of bacterivorous protozoa to their prey allowing pili-producing cells to evade grazing. Further research should define: i) the biophysical differences between pili that allow attachment and floatation, ii) the resources required for this floatation system and advantages over other mechanisms such as flagella, and iii) the ecological trade-offs of having such extracellular appendages. Thus, besides being beneficial, pili could also be a handicap to those cells that produce them as, being phage binding sites^25,26^, they will increase the chance of interacting with phage and hence their susceptibility to phage infection.

The biological carbon pump and microbial loop pose important challenges for streamlined marine cyanobacteria, which have moved away from canonical flagellar motility and have evolved this more passive mechanism for flotation that may require less resources and no additional convoluted tactic systems. This discovery changes our ecological perception of this dominant marine bacterial group, and will have important consequences for our future understanding of predator-prey and carbon flux dynamics in the oceans.

## Methods

### Culture conditions

*Synechococcus* sp. WH7803 was grown in ASW medium and oligotrophic seawater using conditions previously described^22^. Experiments were performed using 20 ml cultures contained in 25 cm² rectangular cell culture flasks (Falcon) with vented caps. Cultures were incubated under optimal growth conditions *i.e.* at 22°C at a light intensity of 10 μmol photons m^−2^ s^−1^ with shaking (140 r.p.m.), unless otherwise stated in the text. To study the influence of nutrients on pili production, media was prepared by i) not adding different nutrient sources into ASW media, *i.e.* nitrogen, phosphorus and trace metals, or ii) diluting ASW in oligotrophic seawater (*i.e.* 1:1000, 1:100 and 1:10). The low light-adapted ecotype *Prochlorococcus* sp. MIT9313 was grown in 40 ml Pro99 medium and PCR-S11 medium with no additional vitamins^33^. Different light intensities (*i.e.* 4 and 15 μmol photons m^−2^ s^−1^) and temperatures (*i.e.* 14 and 22°C) were tested. Cyanobacterial culture growth was routinely monitored by flow cytometry (BD Fortessa).

Grazing experiments were performed using ASW-washed *Uronema marinum* (isolated from Qingdao Bay) and *Cafeteria roenbergensis* CCAP 1900/1 cells. Briefly, 10 ml culture was subjected to centrifugation at 4000g for 15 min and the pellet resuspended in 10 ml ASW. One ml of washed grazers was used to inoculate 20 ml cultures of wild type and pili mutant of *Synechococcus* sp. WH7803 at an initial concentration of 3-4 x 10^6^ cells ml^−1^. Triplicates cultures were incubated under optimal conditions, including shaking to avoid pili mutant sedimentation (see above).

### Pilus knockout mutant in Synechococcus sp. WH7803

Genes SynWH7803_1795 and SynWH7803_1796 (*pilA1* and *pilE*, respectively) were replaced by a gentamicin cassette via a double recombination event to generate the *Synechococcus* sp. WH7803 pili mutant. Two flanking regions of 700 bp from the genome of *Synechococcus* sp. WH7803 and the gentamicin cassette from pBBR-MCS^34^ were amplified by PCR using primers indicated in Table M1, and inserted into vector pK18mobsacB^35^ using the Gibson assembly method following manufacturer’s instructions (New England Biolabs). The detailed protocol to generate mutants in *Synechococcus* sp. WH7803 is given in the Supplementary Information. All three transconjugant colonies that were picked had the same sinking phenotype, had doubly recombined (as checked by sequencing the overlapping regions) and were fully segregated. One mutant was subsequently selected to make axenic by eliminating the ‘helper’ strain and used for further experimentation. The pilus mutation is stable and has retained its sinking phenotype over time.

### Transmission Electron Microscopy

Optimally-grown *Synechococcus* sp. WH7803 and pilus mutant cultures were fixed using 3% (v/v) final concentration glutaraldehyde after which 5 μl were delicately transferred onto a glow-discharge formvar/carbon coated grid and left 2-3 min for cells to attach. After blotting the excess media, negative staining was achieved by applying a drop of 2% uranyl acetate to the grid for 1 min. The excess stain was blotted off and left to air dry before imaging using a JEOL 2011 TEM with Gatan Ultrascan.

### Tracking and imaging of sinking cells

The movement of wild type *Synechococcus* sp. WH7803 and pilus mutant through a nutrient step gradient was performed by placing washed cells in oligotrophic SW on top of a column where nutrient layers (nutrient deplete and amended layers as indicated, using ASW media) were achieved by increasing sucrose concentration (*i.e.* 2.5% w/v per layer).

Sedimentation tracking and velocity measurements were conducted using a setup as previously described^36^. Briefly, a sample chamber was prepared by gluing a square glass capillary (inner dimensions 1.00 x 1.00 mm, length 50 mm; CM Scientific, UK) onto a glass slide using an optical glue (#81; Norland, USA). Tubings (Masterflex Transfer Tubing, Tygon^®^ ND-100-80 Microbore, 0.020" ID x 0.060" OD; inner dimension 0.51 mm; Cole-Parmer, USA) were attached to both ends of the sample chamber using blunt dispensing nozzle tips (Polypro Hubs; Adhesive Dispensing, UK), one-way stopcock valves (WZ-30600-00; Cole-Parmer, USA), and Luer connectors. *Synechococcus* was pulled into the capillary through the inlet by manual suction using a 2 ml syringe connected to the outlet of the capillary system. After introducing the sample, the microfluidic system was isolated by closing the stopcock valves. Samples were left to settle for 30-60 minutes before recording. The slide-capillary system was held vertically by an adjustable translation stage (PT1B/M; Thorlabs, USA), and placed between a white LED ring light source and a continuously focusable objective (InfiniProbe TS-160; Infinity Photo-Optical Company, USA) for dark-field imaging. Images were acquired with magnification set at 16×, using a CMOS FLIR Grasshopper3 (GS3-U3-23S6M-C; Point Grey Research Inc., Canada) operated with FlyCapture2 (FLIR Systems UK). Recordings were done at 1 fps for either 1, 2, or 5 minutes. The system was calibrated with a resolution target (R2L2S1P Positive NBS 1963A; Thorlabs, USA). Sedimentation velocities were calculated with custom-made codes in MATLAB 2019a, based on particle tracking code from Crocker and Grier^37^. Briefly, features corresponding to individual cells were selected based on shape and image intensity, and the distribution of frame-to-frame displacements was calculated from all the tracks lasting at least 20s. All the experiments were checked for outliers.

### Proteome preparation and shotgun analysis

20 ml cell cultures were subjected to centrifugation at 4000g for 15 min at 4°C. Cell pellets were used for cell proteome analyses whereas the supernatants, which were further filtered to remove any remaining cells (0.22 μm), were used for exoproteomic analyses. Exoproteomes were concentrated using a trichloroacetic acid precipitation protocol as previously described^38^. Cell pellets and exoproteome precipitates were resuspended using 1x LDS buffer (ThermoFisher) containing 1% beta-mercaptoethanol and prepared for LC-MS/MS via an in-gel trypsin digestion as described previously^38^. Tryptic peptides were analysed with an Orbitrap Fusion mass spectrometer (Thermo Scientific) coupled to an Ultimate 3000 RSLCnano system (Dionex), using conditions described in Christie-Oleza et al^39^. Mass spectra were identified using *Synechococcus* sp. WH7803 and *Prochlorococcus* sp. MIT9313 UniProt databases (downloaded on 09/11/2017) and quantified using default parameters in MaxQuant v1.6.10.43^40^. Comparative proteomic analyses were performed using MS intensity values in Perseus v1.6.2.2^41^.

### In silico analysis of pilus proteins

Major pilin proteins and pilus machinery were searched in *Synechococcus* sp. WH7803 using reference proteins from *S. elongatus* PCC 7942^16^ and proposed modelled structure^11^. Genomes from cultured picocyanobacteria (*Synechococcus*, n=46, and *Prochlorococcus*, n=43) were downloaded from Cyanorak and re-annotated using PROKKA vs. 1.7^42^. Genomes were then screened for the presence of PilA-like proteins, structural proteins PilQ, PilB and PilC, and competence proteins ComEA and ComEC, via a local BLAST server using the amino acid sequences from *Synechococcus* sp. WH7803.

The 190 picocyanobacterial SAGs from Berube *et al*^28^, all with over 75% estimated genome completeness, were downloaded and annotated in-house using PROKKA vs. 1.7. Annotated SAGs were screened for pilus associated proteins using those from *Synechococcus* sp. WH7803 (Table S1), with further verification using the Conserved Domain search tool from NCBI and manual curation in order to eliminate redundant matches within each SAG and select PilA1-like proteins which had a PilE-like protein encoded in close proximity in the genome.

The PilA1 sequences obtained from the cultured isolates and SAGs were used to generate an HMM profile in Unipro UGENE vs. 33^43^ implemented with the hmmbuild programme from HMM3^44^ using the default parameters. The resulting HMM profile was used to search the TARA oceans metagenomes and metatranscriptomes via the functions offered in the Ocean Gene Atlas portal^29^.

The modelled 3D protein structure of PilA-like proteins was performed with mature sequences using the I-TASSER server’s default settings^45^.

## Acknowledgments

We thank Dr Bakker and the Advanced Bioimaging Platform as well as Dr Hernandez-Fernaud and the WPH Proteomic Facility, at the University of Warwick, for respective support in imaging and proteomics. We thank Dr Guillonneau for providing the protist cultures and WISB centre (grant ref. BB/M017982/1) for access to equipment. This work was supported by NERC grant NE/K009044/1, Ramón y Cajal contract RYC-2017-22452 and the MINECO project CTM2015-70180-R.

## Author contribution

JC-O conceived the study. MA-F generated the knockout mutant. JC-O and MA-F performed the sinking experiments. VZ, AC and DS performed the proteomic analyses. JC-O, RB and RJP performed the bioinformatics analyses. ML and MP carried out the tracking and imaging of sinking cells. JC-O and MA-F wrote the manuscript with large input from DJS and RJP.

## Data and materials availability

All detailed methods and data is available as supplementary information and Tables. The mass spectrometry proteomics data have been deposited in the ProteomeXchange Consortium via the PRIDE partner repository with the dataset identifiers: PXD018394, PXD018395, PXD018396, PXD018524 and PXD019315.

## Supplementary Information

Protocol for generating mutants in marine *Synechococcus*

Extended data Table M1 and Figures S1 to S4

Extended data Tables S1 to S7

Extended data movies 1 to 4

## References

1. Field, C. B., Behrenfeld, M. J., Randerson, J. T. & Falkowski, P. Primary production of the biosphere: integrating terrestrial and oceanic components. Science. 281, 237–240 (1998).

2. Jardillier, L., Zubkov, M. V, Pearman, J. & Scanlan, D. J. Significant CO_2_ fixation by small prymnesiophytes in the subtropical and tropical northeast Atlantic Ocean. ISME J. 4, 1180–1192 (2010).

3. West, N. J. & Scanlan, D. J. Niche-partitioning of *Prochlorococcus* populations in a stratified water column in the eastern North Atlantic Ocean. Appl. Environ. Microbiol. 65, 2585–2591 (1999).

4. Malmstrom, R. R. et al. Temporal dynamics of *Prochlorococcus* ecotypes in the Atlantic and Pacific oceans. ISME J. 4, 1252–1264 (2010).

5. Thompson, A. W. et al. Dynamics of *Prochlorococcus* diversity and photoacclimation during short-term shifts in water column stratification at station ALOHA. Front. Mar. Sci. 5, 1252–1264 (2018).

6. Scanlan, D. J. et al. Ecological genomics of marine picocyanobacteria. Microbiol. Mol. Biol. Rev. 73, 249–299 (2009).

7. Waterbury, J. B., Willey, J. M., Franks, D. G., Valois, F. W. & Watson, S. W. A cyanobacterium capable of swimming motility. Science. 230, 74–76 (1985).

8. McCarren, J. & Brahamsha, B. Transposon mutagenesis in a marine *Synechococcus* strain: isolation of swimming motility mutants. J. Bacteriol. 187, 4457–62 (2005).

9. Cermak, N. et al. Direct single-cell biomass estimates for marine bacteria via Archimedes’ principle. ISME J. 11, 825–828 (2017).

10. Giltner, C. L., Nguyen, Y. & Burrows, L. L. Type IV pilin proteins: versatile molecular modules. Microbiol. Mol. Biol. Rev. 76, 740–772 (2012).

11. Craig, L., Forest, K. T. & Maier, B. Type IV pili: dynamics, biophysics and functional consequences. Nat. Rev. Microbiol. 17, 429–440 (2019).

12. Ellison, C. K. et al. Retraction of DNA-bound type IV competence pili initiates DNA uptake during natural transformation in *Vibrio cholerae*. Nat. Microbiol. 3, 773–780 (2018).

13. Chang, Y.-W. et al. Architecture of the type IVa pilus machine. Science. 351, aad2001 (2016).

14. Bhaya, D., Bianco, N. R., Bryant, D. & Grossman, A. Type IV pilus biogenesis and motility in the cyanobacterium *Synechocystis* sp. PCC6803. Mol. Microbiol. 37, 941–951 (2000).

15. Yoshihara, S. et al. Mutational analysis of genes involved in pilus structure, motility and transformation competency in the unicellular motile ccyanobacterium *Synechocystis* sp. PCC6803. Plant Cell Physiol. 42, 63–73 (2001).

16. Taton, A. et al. The circadian clock and darkness control natural competence in cyanobacteria. Nat. Commun. 11, 1688 (2020).

17. Nagar, E. et al. Type 4 pili are dispensable for biofilm development in the cyanobacterium *Synechococcus elongatus*. Environ. Microbiol. 19, 2862–2872 (2017).

18. Schuergers, N. & Wilde, A. Appendages of the cyanobacterial cell. Life 5, 700–715 (2015).

19. Christie-Oleza, J. A., Armengaud, J., Guerin, P. & Scanlan, D. J. Functional distinctness in the exoproteomes of marine *Synechococcus*. Environ. Microbiol. 17, 3781–3794 (2015).

20. Kaur, A., Hernandez-Fernaud, J. R., Aguilo-Ferretjans, M. D. M., Wellington, E. M. & Christie-Oleza, J. A. 100 Days of marine *Synechococcus*-*Ruegeria pomeroyi* interaction: A detailed analysis of the exoproteome. Environ. Microbiol. 20, 785–799 (2018).

21. Strom, M. S., Nunn, D. N. & Lory, S. A single bifunctional enzyme, PilD, catalyzes cleavage and N-methylation of proteins belonging to the type IV pilin family. Proc. Natl. Acad. Sci. U. S. A. 90, 2404–2408 (1993).

22. Christie-Oleza, J. A., Sousoni, D., Lloyd, M., Armengaud, J. & Scanlan, D. J. Nutrient recycling facilitates long-term stability of marine microbial phototroph-heterotroph interactions. Nat. Microbiol. 2, 17100 (2017).

23. Helaine, S., Dyer, D. H., Nassif, X., Pelicic, V. & Forest, K. T. 3D structure/function analysis of PilX reveals how minor pilins can modulate the virulence properties of type IV pili. Proc. Natl. Acad. Sci. U. S. A. 104, 15888–15893 (2007).

24. Kolappan, S. et al. Structure of the *Neisseria meningitidis* Type IV pilus. Nat. Commun. 7, 13015 (2016).

25. Loeb, T. Isolation of a bacteriophage specific for the F plus and Hfr mating types of *Escherichia coli* K-12. Science. 131, 932–933 (1960).

26. McCutcheon, J. G., Peters, D. L. & Dennis, J. J. Identification and characterization of type IV pili as the cellular receptor of broad host range *Stenotrophomonas maltophilia* bacteriophages DLP1 and DLP2. Viruses 10, E338 (2018).

27. Hu, D. & Reeves, P. R. The remarkable dual-level diversity of prokaryotic flagellins. mSystems 5, (2020).

28. Berube, P. M. et al. Single cell genomes of *Prochlorococcus*, *Synechococcus*, and sympatric microbes from diverse marine environments. Sci. data 5, 180154 (2018).

29. Villar, E. et al. The Ocean Gene Atlas: exploring the biogeography of plankton genes online. Nucleic Acids Res. 46, W289–W295 (2018).

30. Laurenceau, R. et al. Toward a genetic system in the marine cyanobacterium *Prochlorococcus*. Access Microbiol. 2, (2020).

31. Coe, A. et al. Survival of *Prochlorococcus* in extended darkness. Limnol. Oceanogr. 61, 1375–1388 (2016).

32. Stocker, R. & Seymour, J. R. Ecology and physics of bacterial chemotaxis in the ocean. Microbiol. Mol. Biol. Rev. 76, 792–812 (2012).

33. Moore, L. R. et al. Culturing the marine cyanobacterium *Prochlorococcus*. Limnol. Oceanogr. Methods 5, 353–362 (2007).

34. Kovach, M. E. et al. Four new derivatives of the broad-host-range cloning vector pBBR1MCS, carrying different antibiotic-resistance cassettes. Gene 166, 175–176 (1995).

35. Kvitko, B. H. & Collmer, A. Construction of *Pseudomonas syringae* pv. tomato DC3000 mutant and polymutant strains. Methods Mol. Biol. 712, 109–128 (2011).

36. Font-Muñoz, J. S. et al. Collective sinking promotes selective cell pairing in planktonic pennate diatoms. Proc. Natl. Acad. Sci. 116, 15997–16002 (2019).

37. Crocker, J. C. & Grier, D. G. Methods of digital video microscopy for colloidal studies. J. Colloid Interface Sci. 179, 298–310 (1996).

38. Christie-Oleza, J. A. & Armengaud, J. In-depth analysis of exoproteomes from marine bacteria by shotgun liquid chromatography-tandem mass spectrometry: the *Ruegeria pomeroyi* DSS-3 case-study. Mar. Drugs 8, 2223–2239 (2010).

39. Christie-Oleza, J. A., Scanlan, D. J. & Armengaud, J. ‘You produce while I clean up’, a strategy revealed by exoproteomics during *Synechococcus*-*Roseobacter* interactions. Proteomics 15, 3454–3462 (2015).

40. Cox, J. & Mann, M. MaxQuant enables high peptide identification rates, individualized p.p.b.-range mass accuracies and proteome-wide protein quantification. Nat. Biotechnol. 26, 1367–1372 (2008).

41. Tyanova, S. et al. The Perseus computational platform for comprehensive analysis of (prote)omics data. Nat. Methods 13, 731–740 (2016).

42. Seemann, T. Prokka: rapid prokaryotic genome annotation. Bioinformatics 30, 2068–2069 (2014).

43. Okonechnikov, K., Golosova, O. & Fursov, M. Unipro UGENE: a unified bioinformatics toolkit. Bioinformatics 28, 1166–1167 (2012).

44. Mistry, J., Finn, R. D., Eddy, S. R., Bateman, A. & Punta, M. Challenges in homology search: HMMER3 and convergent evolution of coiled-coil regions. Nucleic Acids Res. 41, e121–e121 (2013).

45. Yang, J. & Zhang, Y. Protein structure and function prediction using I‐TASSER. Curr. Protoc. Bioinforma. 52, 5.8.1–5.8.15 (2015).

