## Supplementary Information for "Pili allow dominant marine cyanobacteria to avoid sinking and evade predation" (from another version of this preprint: version 1, 2020-07-09)

**This file includes**

Protocol for generating mutants in marine *Synechococcus*

Extended data table M1 and figures S1 to S4

**Protocol for generating mutants in marine *Synechococcus***

This protocol is largely based on the one published by Brahamsha[1] but with the difference of adding an engineered *Ruegeria pomeroyi* DSS-3 strain that can later be eliminated to regenerate axenic *Synechococcus* cultures. The conjugation-recombination-plating efficiency of the process is below ~$10^{-9}$-$10^{-10}$ and, hence, each mutant is amongst the debris of $10^{9}$-$10^{10}$ dead cells which we know is toxic for these marine cyanobacteria[2]. *R. pomeroyi* DSS-3 functions as a helper strain to detoxify the cell debris generated during the process[3] and enhances the survival of mutant cells. *R. pomeroyi* DSS-3 contains the fusion plasmid pKNG101-BBR-MCS (see below for further details) which confers, in the first instance, resistance to the chosen antibiotic (*e.g.* Km or Gm), and then allows the elimination of the heterotroph with sucrose as it carries a lethal *sacB* gene.

This method has been successfully tested with *Synechococcus* strains WH7803 and WH8102. In all cases, the use of the helper strain *R. pomeroyi* DSS-3 generated tens of *Synechococcus* mutant colonies per conjugation whereas no colonies were obtained when the heterotroph was not included.

*Plasmid constructs*

- Most plasmids are unable to replicate in marine *Synechococcus*. Hence, most vectors can be used to generate mutants in these cyanobacteria. We used the conjugative plasmid pK18mobsacB[4], although we have also successfully tested pGP704[5] in which we had inserted the Km resistance cassette from pBBR-MCS1Km[6].
- All *Synechococcus* strains tested (*i.e.* WH7803 and WH8102) are sensitive to Km and Gm, and can become resistant to these antibiotics through the insertion of a resistance gene cassette. This is not the case for Cm, where resistance was not achieved despite the correct insertion of a resistance cassette.
- We recommend double crossover mutations (*i.e.* replacing the gene of interest for a marker gene *e.g.* Gm resistance gene). Single crossovers, in which the gene of interest is disrupted through the insertion of the whole plasmid, have been successfully generated although the integration of the plasmid into the genome of *Synechococcus* seems to generate problems of instability (*i.e.* reversion may occur when antibiotic pressure is not maintained).
- Four-fragment plasmid constructs as shown in the diagram below can be easily generated using methods such as the Gibson assembly kit.

*Conjugative E. coli strain*

- The plasmid is delivered to *Synechococcus* via conjugation. We recommend the use of *E. coli* S17.1λpir[7] as it contains all the machinery necessary for conjugation stably integrated in its chromosome.

*Helper strain*

- *Ruegeria pomeroyi* DSS-3 is clearly a good helper strain for marine *Synechococcus*[2]. It helps the growth of *Synechococcus* through the breakdown of leaked or dead-cell debris which otherwise becomes toxic to the mutant cells inhibiting their growth. The addition of this helper strain makes a clear difference and, in it, resides the novelty of this protocol.
- Our helper strain *R. pomeroyi* DSS-3 contains plasmid pKNG101-BBR-MCS which comes from the fusion of plasmids pKNG101[8] and pBBR-MCS1Gm[6]. Hence, it confers resistance to Gm (or other

antibiotics of choice, *e.g.* Km) and contains the lethal gene *sacB* so it can be eliminated at the end of the process to redeem an axenic mutant *Synechococcus* culture.

*Materials*

- ASW medium[9] is routinely used to grow marine *Synechococcus*.
- One plate of ASW agarose 0.3% (w/v) as the platform for conjugation. Let it dry so it will absorb the liquid from the conjugation.
- Molten ASW agarose 0.2% (w/v) brought to 37°C to plate the conjugation.

*Protocol*

- Grow a 100 ml *Synechococcus* culture in optimal conditions to high cell densities (> $10^8$ cells $ml^{-1}$; ~2 weeks), in ASW medium, 22°C at a light intensity of 10-15 µmol photons $m^{-2}$ $s^{-1}$ with shaking (140 rpm).
- Grow *E. coli* S17.1λpir containing the plasmid construct overnight in 10 ml LB with no antibiotics. Incubate at 30°C with shaking (140 rpm). Plasmids are unstable in this *E. coli* strain when incubated at higher temperatures (do NOT grow over 30°C!).
- Harvest the *Synechococcus* and *E. coli* cells by centrifugation (3,200 g at 23°C for 15 min).
- Discard the supernatant and resuspend the pellet in the small volume of remaining media. Mix the *Synechococcus* and *E. coli* cells and transfer to an Eppendorf.
- Pellet the cells by centrifugation (13,000 g at 23°C for 1 min) and resuspend in ~200µl of remaining supernatant.
- Place the cells on an ASW agarose 0.3% plate and leave to dry until the liquid has been absorbed by the plate. Alternatively, the conjugation can be placed on a 0.2 µm pore size filter to facilitate the cell collection.
- Parafilm the plate and leave it upright. Incubate for 48h at 23°C and a light intensity of 10 µmol photons $m^{-2}$ $s^{-1}$.
- Grow the helper strain *R. pomeroyi* DSS-3 (containing the plasmid pKNG101-BBR-MCS with the correct antibiotic cassette) in Marine Broth and incubate for 48h at 28°C with shaking (140 rpm). 3 ml of culture is enough.
- After 48h, pellet 2 ml of the *R. pomeroyi* culture by centrifugation (13,000 g at 23°C for 1 min) and resuspend the cells in 1 ml ASW.
- Collect the *Synechococcus*-*E. coli* conjugation by pipetting in 1 ml ASW.
- Cool 100 ml of molten ASW agarose 0.2% (w/v) to 37°C and add: i) the *R. pomeroyi* cells (diluting 1:100, achieving a final concentration ~$10^6$-$10^7$ cells $ml^{-1}$), and ii) the antibiotic (e.g. Km 50 µg $ml^{-1}$ or Gm 15 µg $ml^{-1}$).
- Dispense 30 ml of this molten ASW agarose in 50-ml falcon tubes. Immediately add *e.g.* 50, 100 or 500 µl of the resuspended conjugation, and pour immediately into a petri dish. Leave to cool/solidify.
- Delicately parafilm the plates and incubate them upright in plastic bags at 23°C and at a light intensity of 5 µmol photons $m^{-2}$ $s^{-1}$. Four to six weeks are required before visible colonies are obtained.
- Disturb small colonies with a sterile pipet tip and leave 4-5 days to grow larger colonies.
- Pipette the colonies out of the petri dishes into 5 ml ASW with the selection antibiotic and incubate under optimal conditions.

- As the culture grows denser, increase the volume by adding ASW plus antibiotic.
- Check by PCR to verify for the correct mutant (*i.e.* the gene of interest has been disrupted) and, in polyploid strains, that the mutant has segregated.
- Take 8 ml of the culture and add 2 ml ASW containing 50% (w/v) sucrose and incubate under optimal conditions for 24h. This step will eliminate most of the *R. pomeroyi* helper strain (which contains *sacB*).
- After 24 h, prepare molten ASW agarose 0.2% (w/v) and cool to 37°C. Add the corresponding antibiotic.
- Dilute the treated culture to extinction and plate 100 µl of the $10^{-5}$, $10^{-6}$ and $10^{-7}$ dilutions as done previously (*i.e.* pipetting the cells into 30 ml of cooled molten ASW agarose 0.2% and immediately plate into Petri dishes).
- Parafilm the plates and incubate them upright in plastic bags at 23°C and a light intensity of 5 µmol photons $m^{-2}$ $s^{-1}$. Four to six weeks are required before visible colonies are obtained.
- Carefully transfer colonies to a new ASW agarose 0.2% plate plus antibiotics and disturb them so they grow larger colonies. Do not disturb colonies in the original plate as you will increase the risk of contaminating the *Synechococcus* colony with the heterotrophic helper that may have survived the sucrose treatment. Be very careful when picking the colonies out of the agar plate to avoid contamination.
- After 4-5 days, check all picked *Synechococcus* colonies for contamination on Marine Agar plates.
- Pipette axenic colonies out of the petri dish into 5 ml ASW with the selection antibiotic and incubate in optimal conditions.

**References**

1. Brahamsha, B. A genetic manipulation system for oceanic cyanobacteria of the genus *Synechococcus*. *Appl. Environ. Microbiol.* **62**, 1747–1751 (1996).
2. Christie-Oleza, J. A., Sousoni, D., Lloyd, M., Armengaud, J. & Scanlan, D. J. Nutrient recycling facilitates long-term stability of marine microbial phototroph-heterotroph interactions. *Nat. Microbiol.* **2**, 17100 (2017).
3. Kaur, A., Hernandez-Fernaud, J. R., Aguilo-Ferretjans, M. D. M., Wellington, E. M. & Christie-Oleza, J. A. 100 days of marine *Synechococcus-Ruegeria pomeroyi* interaction: A detailed analysis of the exoproteome. *Environ. Microbiol.* **20**, 785–799 (2018).
4. Kvitko, B. H. & Collmer, A. Construction of *Pseudomonas syringae* pv. tomato DC3000 mutant and polymutant strains in *Plant Immunity. Methods in Molecular Biology* 109–128 (2011).
5. Miller, V. L. & Mekalanos, J. J. A novel suicide vector and its use in construction of insertion mutations: osmoregulation of outer membrane proteins and virulence determinants in *Vibrio cholerae* requires *toxR*. *J. Bacteriol.* **170**, 2575–2583 (1988).
6. Kovach, M. E. *et al.* Four new derivatives of the broad-host-range cloning vector pBBR1MCS, carrying different antibiotic-resistance cassettes. *Gene* **166**, 175–176 (1995).
7. Herrero, M., de Lorenzo, V. & Timmis, K. N. Transposon vectors containing non-antibiotic resistance selection markers for cloning and stable chromosomal insertion of foreign genes in gram-negative bacteria. *J. Bacteriol.* **172**, 6557–6567 (1990).
8. Kaniga, K., Delor, I. & Cornelis, G. R. A wide-host-range suicide vector for improving reverse genetics in Gram-negative bacteria: inactivation of the *blaA* gene of *Yersinia enterocolitica*. *Gene* **109**, 137–141 (1991).

9. Wilson, W. H., Carr, N. G. & Mann, N. H. The effect of phosphate status on the kinetics of cyanophage infection in the oceanic cyanobacterium *Synechococcus* sp. WH7803. *J. Phycol.* **32**, 506–516 (1996).

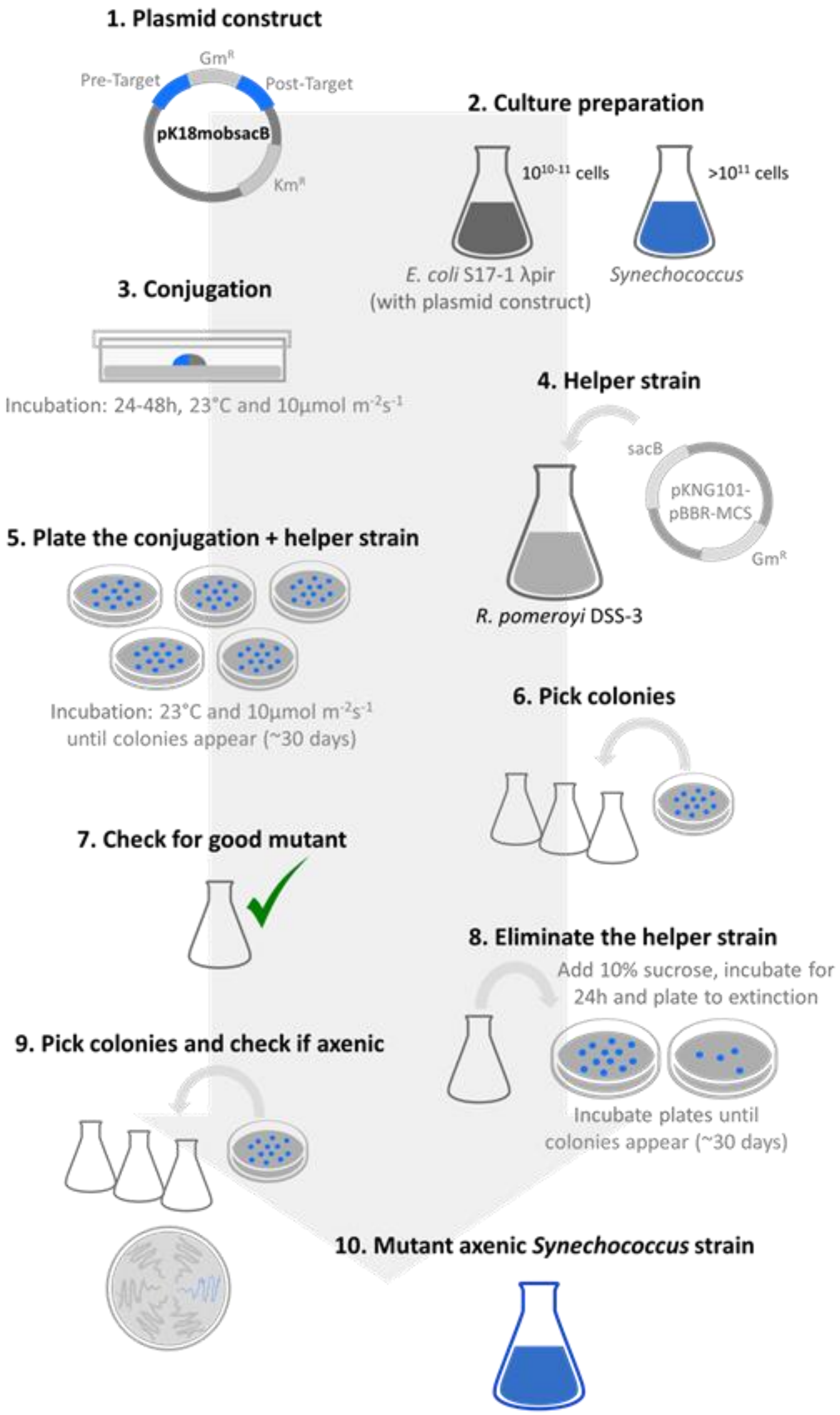

### Supplementary Tables and Figures

**Table M1.** Primers for PCR amplification of the fragments used to assemble the vector to generate the pili knockout mutant.

| Primer name | Sequence | Annealing temperature |
|---|---|---|
| Pili_pre_fwd | gccacctgcagatctgcaggCCAGAAGCAGCTGTTCATCGG | 3'Tm=66.7 3'Ta(annealing temp)=69.7 |
| Pili_pre_rev | cattcatcccTGGCTGGTCCGTGGATTGGG | 3'Tm=71.0 3'Ta(annealing temp)=69.7 |
| Km_fwd | ggaccagccaGGGATGAATGTCAGCTACTG | 3'Tm=60.4 3'Ta(annealing temp)=61.2 |
| Km_rev | attgaattttTGCATAAAAACTGTTGTAATTCATTAAG | 3'Tm=58.2 3'Ta(annealing temp)=61.2 |
| Pili_post_fwd | tttttatgcaAAAAATTCAATACAATGAAACCATTC | 3'Tm=56.3 3'Ta(annealing temp)=59.3 |
| Pili_post_rev | tcaaaacaaactagcgatcgGTTATCAGTATCGTTTGGCATC | 3'Tm=59.0 3'Ta(annealing temp)=59.3 |

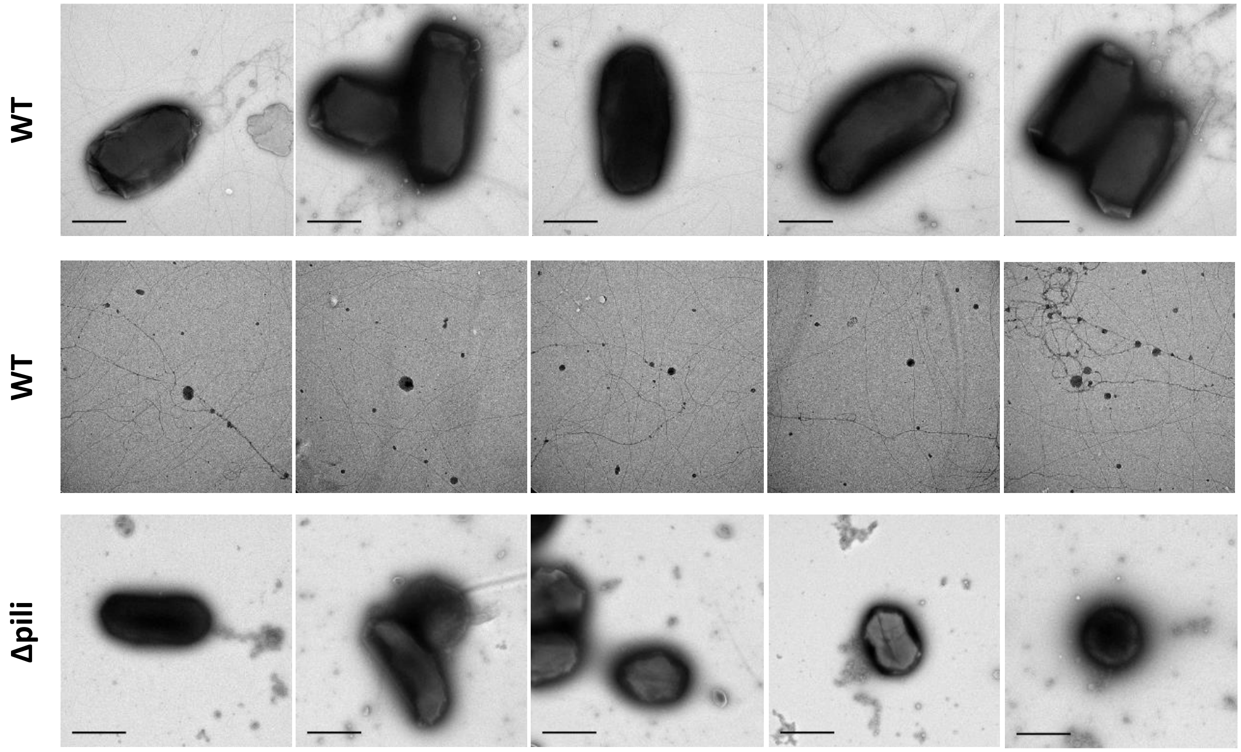


**Fig S1.** Transmission electron microscope images of pili structures in wild type (WT) *Synechococcus* sp. WH7803, and their absence in the pili mutant (Δpili). Middle panel images, obtained with the same magnification as other panels, represent intercellular regions of wild type cells to improve the visualisation of the pili. Scale bar represents 1 µm. Cells were obtained from late exponential phase liquid cultures incubated in ASW medium under optimal growth conditions (including shaking).

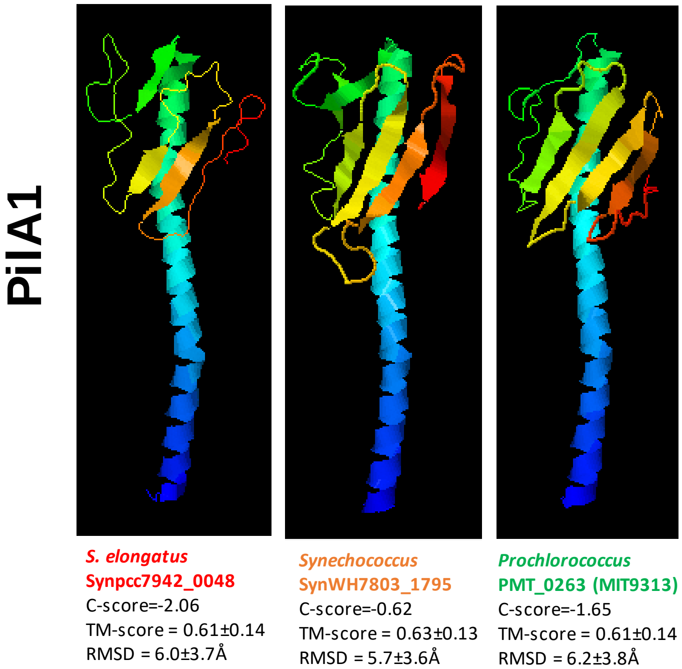


**Fig S2.** Modelled 3D protein structure of PilA1, PilA2, PilA3 and PilA from different marine bacterial strains, *Synechococcus elongatus* PCC 7942 and pathogens. Models were performed using the server I-TASSER with amino acid sequences where the signal peptide had been removed.

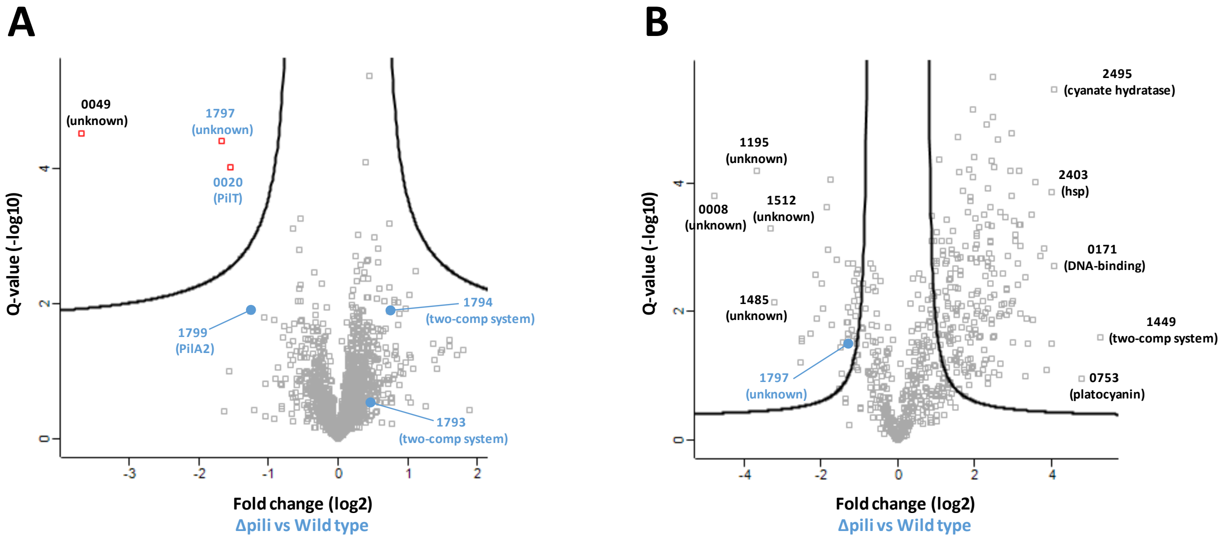


**Fig S3.** Volcano plots obtained from comparative cellular (A) and extracellular (B) proteomic analysis between wild type *Synechococcus* sp. WH7803 and pili mutant (Δpili). Proteins related to the pilus are highlighted in blue. While the cellular proteome showed a small variance between proteomes, the exoproteomes were highly divergent despite efforts to normalise the loss of the highly abundant PilA1 in the pili mutant. This was mainly caused by the detection of low abundance proteins in the exoproteomes of Δpili which may have been masked and remained undetected in the wild type strain due to the presence of PilA1.

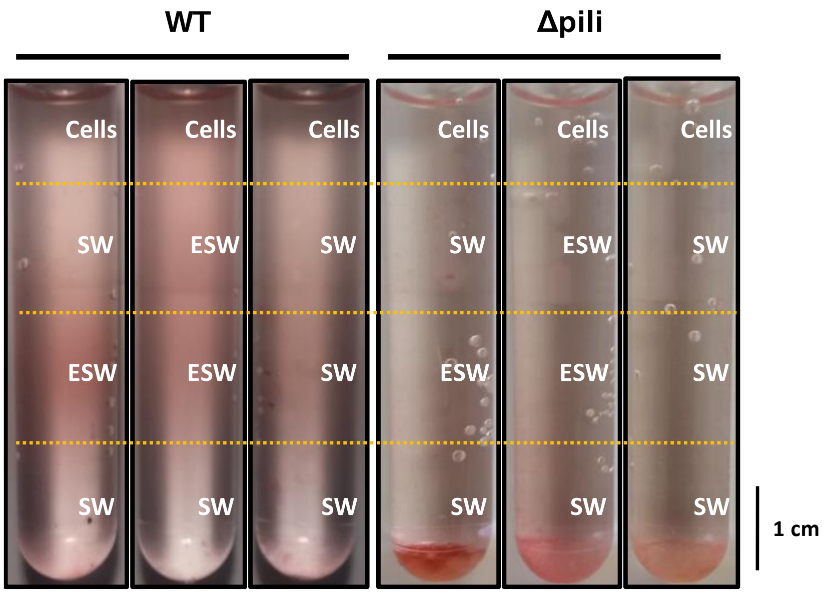


**Fig S4.** Influence of the pili in the positioning of cells within the water column. Nutrient step gradient column where wild type and pili mutant (Δpili) cells of *Synechococcus* sp. WH7803 were placed at the top and incubated for three days. Nutrient deplete (SW) and nutrient enriched layers (ESW) are indicated.
